# Chemical protein unfolding - A simple cooperative model

**DOI:** 10.1101/2023.05.11.540313

**Authors:** Joachim Seelig, Anna Seelig

## Abstract

Chemical unfolding with guanidineHCl or urea is a common method to study the conformational stability of proteins. The analysis of unfolding isotherms is usually performed with an empirical linear extrapolation method (LEM). The method contradicts however, common expectations. A large positive free energy is assigned to the native protein which is usually considered to be a minimum of the free energy. Here we present a multistate cooperative model, which addresses specifically the binding of the denaturant to the protein and the cooperativity of the protein unfolding equilibrium. The model is based on a molecular statistical-mechanical partition function of the ensemble but simple solutions for the calculation of the binding isotherm and the associated free energy are presented. The model is applied to 23 published unfolding isotherms of small and large proteins. For a given denaturant, the binding constant depends on temperature and pH, but shows little protein specificity. Chemical unfolding is less cooperative than thermal unfolding. The cooperativity parameter σ is two orders of magnitude larger than that of thermal unfolding. The multistate cooperative model predicts a zero free energy for the native protein, which becomes strongly negative beyond the midpoint concentration of unfolding. The free energy to unfold a cooperative unit corresponds exactly to the diffusive energy of the denaturant concentration gradient necessary for unfolding. The temperature dependence of unfolding isotherms yields the denaturant-induced unfolding entropy and, in turn, the unfolding enthalpy. The multistate cooperative model provides molecular insight, is as simple to apply as the LEM, but avoids the conceptual difficulties of the latter.

## Introduction

Chemical denaturation is a process by which the protein conformation is unfolded via addition of denaturants such as guanidineHCl, urea or SDS. Chemical denaturation is a common method for determining a protein’s conformational stability, relative to its functional properties.^1, 2^ Identifying the conditions that maximize the structural stability of a protein is crucial during development of biologics for therapeutic treatments. Several complementary techniques should be applied to provide a systematic analysis of protein stability.^3^

Chemical unfolding of proteins is analysed almost exclusively with a 2-state model, the Linear Extrapolation Method (LEM).^4^ The LEM is an empirical approximation and its conceptual difficulties have been realized since its initial proposal.^4–6^ In particular, the LEM assigns a large positive Gibbs free energy 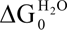 to the native protein. However, “the general understanding in the protein folding field has been that proteins fold into their native conformations driven by decrease in Gibbs free energy (negative Δ*G*).”^7^ This thermodynamic hypothesis has become the default physical description of protein folding. In this view the native state is the most stable one, that is, the global *G* minimum, not a maximum.

The 2-state model is a non-cooperative model. It has no energy parameter for the interaction between neighboring amino acid residues. All amino acid residues unfold simultaneously (all-or-none model).^8^ However, the conformational change of all amino acid residues at the same time is physically unrealistic. Instead, “peptides that form helices in solution do not show a simple two-state equilibrium between a fully folded and fully unfolded structure. Instead they form a complex mixture of all helix, all coil or, most frequently central helices with frayed coil ends”.^9^ A sequential cooperative unfolding of protein domains is therefore a physically more realistic alternative.

We have recently proposed a semi-empirical model that describes a cooperative protein unfolding.^10^ The model assumes explicitly the binding of the denaturant D to the protein with the binding constant K_D_ and the cooperative unfolding of the protein with the cooperativity parameter σ ^10^ Here we provide a modification of this model, based on a statistical-mechanical partition function leading to simple expressions for the chemical unfolding isotherm and the associated free energy.

Published chemical unfolding isotherms obtained with spectroscopic techniques and with calorimetry are analyzed. GuanidineHCl, urea and sodium dodecyl sulfate (SDS) were studied as denaturants. The protein size ranged from 30 to ∼1600 amino acid residues, including a monoclonal antibody. The present model yields excellent simulations of all unfolding isotherms. The native protein is the reference state with zero free energy. The free energy becomes negative upon unfolding and decreases with the logarithm of the denaturant concentration. The temperature dependence of the free energy provides the unfolding entropy and, in turn, the unfolding enthalpy. The latter agrees with calorimetric measurements. The binding constant K_D_ depends essentially on the type of denaturant and varies little with the nature of the protein. The cooperativity parameter for chemical unfolding is compared to that for thermal unfolding.^11^ The multistate cooperative model is firmly grounded on statistical mechanical thermodynamics. With a multistate cooperative approximation, we provide a simple expression for cooperative chemical denaturation analysis which is equally easy to apply as the LEM.

## Materials and Methods

Chemical unfolding experiments with guanidine HCl, urea, and sodium dodecyl sulfate SDS), performed with different spectroscopic and calorimetric techniques, are selected from the available literature. The focus is on the chemical unfolding of lysozyme with guanidineHCl, urea, and SDS. Altogether 23 published chemical denaturation isotherms of proteins of different structure and size have been investigated.

### Chemical unfolding models

Chemical denaturants such as guanidineHCl or urea are commonly used to study protein stability. They change the polarity of the environment, bind to backbone and amino acid residues and thus change the native protein conformation (N) into the unfolded conformation (U).

#### Multistate cooperative unfolding model

The multistate cooperative model considers the individual amino acid residues of the protein. The amino acid residues in the native or in the unstructured confirmation are designated as “n” and “u”. The initial step of this model is the binding of a denaturant D to an amino acid “n” in a structured protein segment, inducing a conformational transition to an unstructured state “u”:

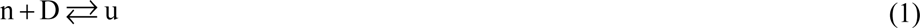

This chemical equilibrium is described by a simple equation:

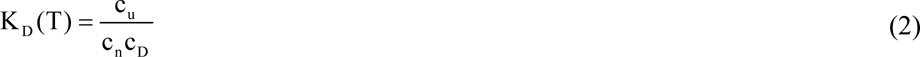

The concentrations c_n_ and c_u_ are the concentrations of the amino acid residues participating in unfolding. The binding constant K_D_(T) is a function of temperature only. Unfolding is a dynamic equilibrium between many different protein conformations.

The statistical interpretation of a chemical equilibrium requires a grand canonical partition function. To make the connection to the textbook literature we consider a system of only two types of particles. “In a grand canonical ensemble, the number of type A particles, N_A_, and type B particles, N_B_, are both variable. Let μ_A_ and μ_B_ be the respective chemical potentials of the two components. The grand partition function is

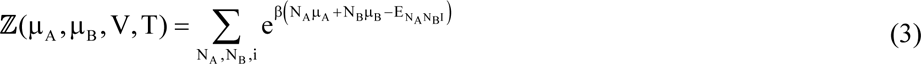

(reference^12^, chapter 13.9). Equation 2 defines a 3 component system measuring the ensemble size in concentration units c_i_ (mol/L). The chemical potential is given by

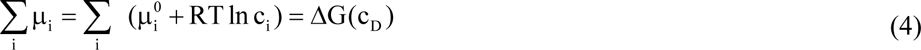

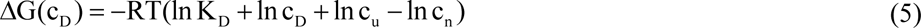

In chemical unfolding experiments the protein concentration is typically ∼ 10 μM, whereas the concentration of the denaturant is 1 - 8 M. The chemical potential of the equilibrium is dominated by the denaturant concentration, leading to the following approximation.

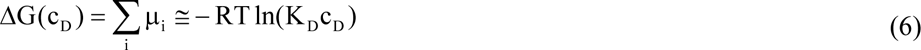

The conditional probability, q(c_D_) of a residue “n” in a structured protein segment is defined as 1, the conditional probability of the unfolded residue “u” is

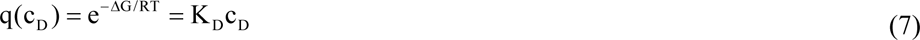

The conditional probability q(c_D_) is inserted into the Zimm-Bragg partition function Z(c_D_)^12^.

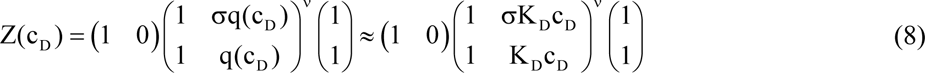

ν denotes the number of amino acid residues participating in the unfolding reaction. The cooperativity parameter σ determines the steepness of the unfolding transition. A small σ leads to a sharp transition. The cooperativity parameter in chemical unfolding is typically σ ∼ 10^-2^ – 10^-3^ and is 10 to hundred times larger than that of thermal unfolding. The fraction of unfolded protein is:

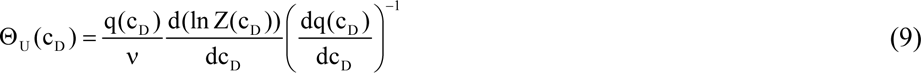

For a non-cooperative ensemble with σ = 1 the partition function eq. 8 reduces to Z(q) = (1+ q)^ν^ . The fraction of unfolded protein becomes independent of ν and is 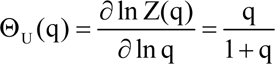, which is identical to the Langmuir adsorption isotherm.

Figure 1 shows the unfolding isotherm Θ_U_(c_D_) (eq. 9) for different cooperativity parameters σ. The binding constant is K_D_ = 0.25 M^-1^, which is typical for guanidineHCl binding to most proteins. The binding constant is too small to induce complete protein unfolding for a non-cooperative ensemble (σ = 1) as demonstrated by the Langmuir isotherm in figure 1A. A dramatic change in the binding isotherm is induced by including even small cooperative interactions (red to green lines in fig. 1A).

**Figure 1.**
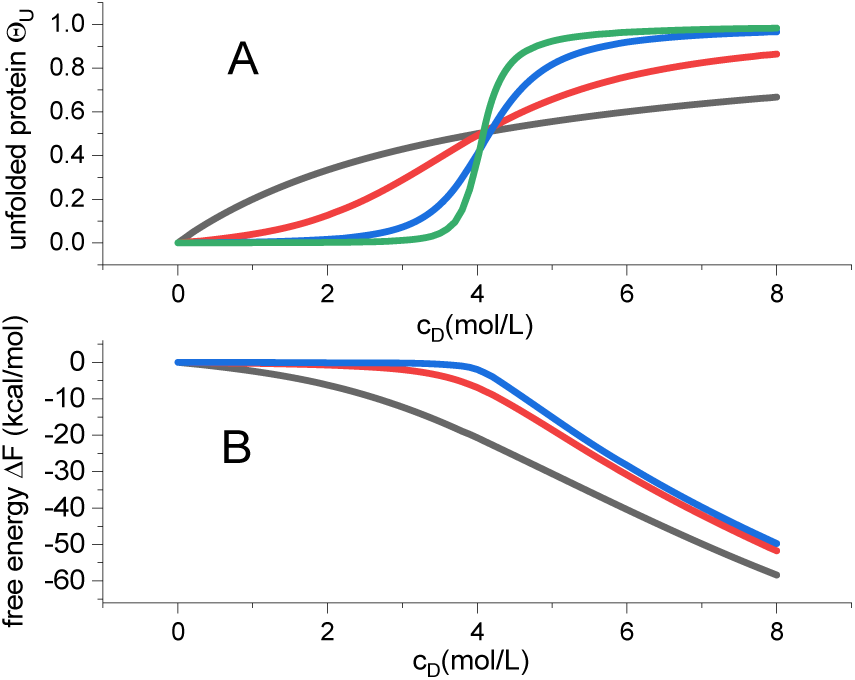
Multistate cooperative unfolding model. Variation of the cooperativity parameter σ. (A) Extent of unfolding Θ_U_(c_D_) (eq. 9). Black: σ = 1. Red: σ = 10^-1^. Blue: σ = 10^-2^. Green: σ = 10^-3^ (B) Free energy as a function of cooperativity parameter σ and denaturant concentration (eq. 10). Binding constant K_D_ = 0.25. M^-1^. Midpoint concentration c_0_ = 4.0 M. Number of amino acid residues participating in transition ν =129. Temperature T=298K. The simulation parameters K_D_, c_0_, ν, T and σ = 10^-3^ correspond to the unfolding of lysozyme in guanidineHCl solution.

The extent of unfolding (eq. 9) is the result of the partition function Z(c_D_). Likewise, the free energy is also related to the partition function according to standard statistical thermodynamics.^13^ The free energy change of unfolding is

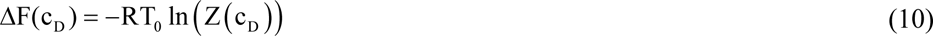

ΔF(c_D_) upon addition of guanidineHCl is displayed in figure 1B. The native protein is the reference state with zero free energy. Upon addition of denaturant the free energy decreases. In the case of no cooperativity (σ = 1), the free energy decreases already at low concentrations of denaturant. For a cooperative ensemble, the free energy remains close to zero up to the midpoint of unfolding c_0_ and decreases rapidly beyond this concentration. Compared to non-cooperative denaturation the free energy change for a co-operative system is distinctly smaller.

The temperature dependence of the free energy is

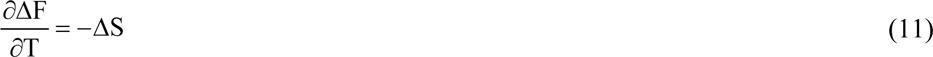

The unfolding enthalpy can then be calculated as

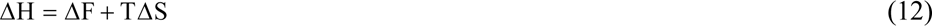

#### A simple multistate cooperative approximation

For the biochemical practitioner the above formalism may act as a deterrent. Fortunately, the matrix equation 7 can be replaced by a simple linear expression, which can easily be calculated. The cooperativity parameter in chemical unfolding experiments is always σ ≥ 10^−3^ and the largest eigenvalue λ_0_ of the above matrix is a sufficient approximation, resulting in a simpler partition function ^12^

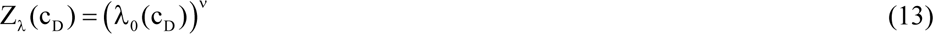

With

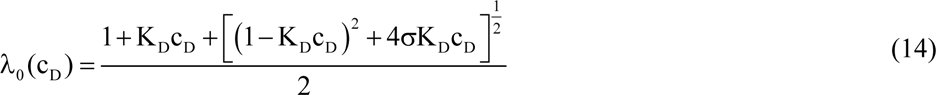

The fraction of unfolded protein is given by

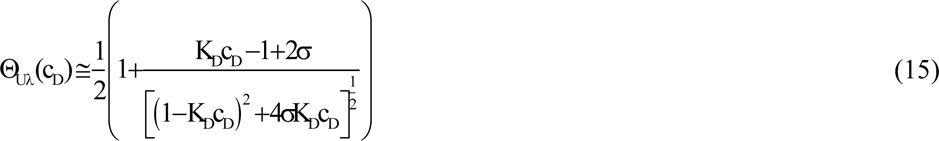

At the midpoint of unfolding with c_D_ = c_0_ and Θ(c_0_) = 1/2 follows

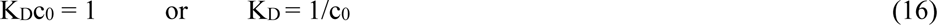

In the multistate cooperative model, the binding constant of the denaturant is simply the reciprocal of the midpoint concentration of unfolding. Equation 15 can also be written as

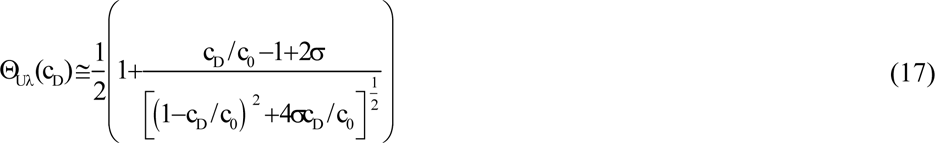

Equations 15 and 17 are equivalent to a cooperative sorption isotherm, which is based on a statistical-mechanical partition function. They are In line with many other sorption isotherms.^14^ The only fit parameter in the simulation of chemical unfolding isotherms is therefore the cooperativity parameter σ.

The free energy of the system is

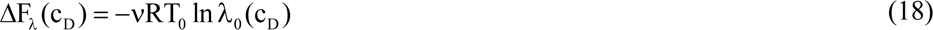

For denaturant concentrations c_D_ ≫ c_0_, the following approximation is valid

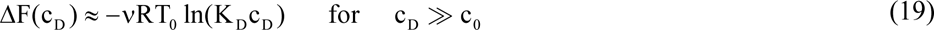

#### *Concentration gradient* Δ*c* = *c_end_* − *c_ini_ and the free energy of the n* → *u transition*

The multistate cooperative model predicts a one-to-one relationship between the free energy of the concentration gradient Δc = c_end_ − c_ini_ and the energy required to induce the n → u transition. Unfolding takes place in the concentration interval c_ini_ ≤ c_D_ ≤ c_end_ . The concentration gradient Δc = c_end_ − c_ini_ constitutes a diffusive or osmotic free energy

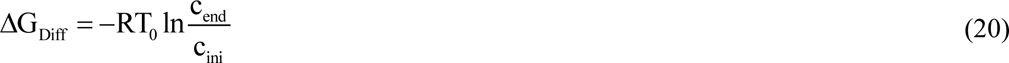

The free energy at ambient temperature is typically ΔG_Diff_ ∼ −0.4 to −1.0 kcal/mol. In parallel, the binding of the D is associated with the free energy of the n → u transition per amino acid residue Δg_nu_ .

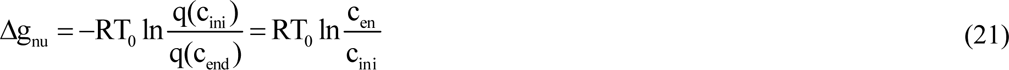

The predicted unfolding free energy per amino acid residue is thus identical to the concentration gradient (with opposite sign), that is, Δg_nu_ + ΔG_diff_ = 0 . The multistate cooperative model is thus consistent with the thermodynamic expectation for a reversible equilibrium.

#### Chemical equilibrium two-state unfolding

The common model to describe chemical unfolding isotherms is a non-cooperative two-state model, which has dominated the field for the last 30 - 40 years. Only two types of protein conformation are assumed in solution, the native protein (N) and the unfolded protein (U). No intermediate structures and no specific interaction between denaturant D and protein are considered. The equilibrium N ⇆ U is simply

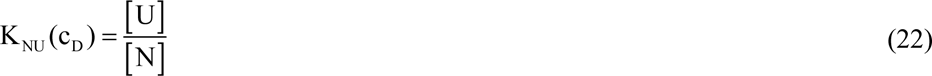

The equilibrium constant K(c_D_) varies with the concentration of denaturant. To calculate K_NU_(c_D_) the model makes an unconventional assumption about the free energy.

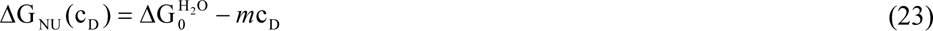

Eq. 21 is difficult to understand in thermodynamic terms for two reasons. First, a large positive free energy 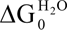 is assigned to the stable native protein. Second, the free energy is a *linear*, not as usual, a logarithmic function of the concentration c_D_. The so-called Linear Extrapolation Method allows the calculation of spectroscopic unfolding transitions. The fraction of unfolded protein Θ_U_(c_D_) is

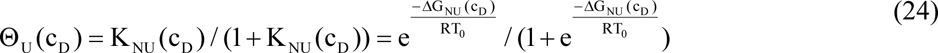

The disadvantage of this model is its conceptual simplicity of just two protein conformations and the rather questionable Linear Extrapolation Method. It is generally believed that the native protein sits in a free energy minimum. In contrast, the LEM predicts a large positive free energy for a stable native protein.

By definition the midpoint concentration c_0_ is the concentration where native and unfolded protein occur at equal concentrations and the free energy is consequently a zero in the LEM. It thus follows that

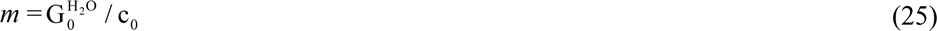

*m* is not an independent variable, which is however often ignored in the analysis of spectroscopic data.

## Results

### Chemical unfolding of lysozyme with guanidineHCl at 30 °C

Figure 2 displays the chemical unfolding of lysozyme in guanidine HCl at 30 °C experimental (data from reference.^15^) The midpoint concentration of unfolding is c_0_ = 3.9 M. Chemical unfolding takes place in the concentration range of. 2.9 M ≤ c_D_ ≤ 5.3 M

**Figure 2.**
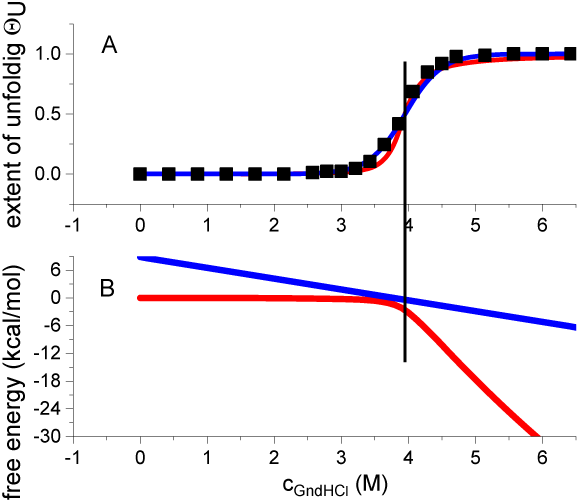
Unfolding of lysozyme in guanidineHCl solution at 30 °C. Red lines: multistate co-operative model. Blue lines: Linear Extrapolation Method. Vertical lines: midpoint concentration c_0_ = 3.9 M (A) (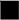) data taken from reference.^15^. Simulation parameters of multistate cooperative mode: K_D_ = 1/c_0_= 0.26 M^-1^, σ = 1×10^−3^ . LEM parameters: 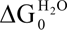 =8.837 kcal/mol, m = 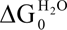 / c =2.34 kcalL/mol^2^. (B) Temperature profiles of the free energy.

The solid lines in figure 2A simulate the experimental data with the parameters given in the legend to figure 2. Both methods simulate the unfolding transitions equally well. Both models predict exactly 50% unfolding at the midpoint concentration c_0,_ but the concentration profiles of the free energy are quite different (figure 2B). The LEM assigns a large positive free energy 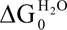 = 8.93 kcal/mol to the native protein. At the midpoint concentration c_0_ the free energy is exactly zero (ΔG(c_0_) = 0). The multistate cooperative model predicts a zero free energy for the native protein. The free energy change is slightly negative up to c_0_, is ΔF = −2.0 kcal/mol at c_0_, and decreases rapidly at c_D_ > c_0_ (eq.18). The change in free energy upon unfolding (in the concentration range 2.9 M ≤ c_D_ ≤ 5.3 M is ΔF = −21.7 kcal/mol for the multistate cooperative model, but only ΔG = −11.8 kcal/mol for the LEM.

### Lysozyme unfolding in urea

Urea is less effective in chemical denaturation than guanidineHCl as is obvious in figure 3. The urea midpoint concentration is 6.7 M compared to 4.1 M for guanidine HCl. All thermodynamic data are summarised in table 2, see ref.^16^

**Figure 3.**
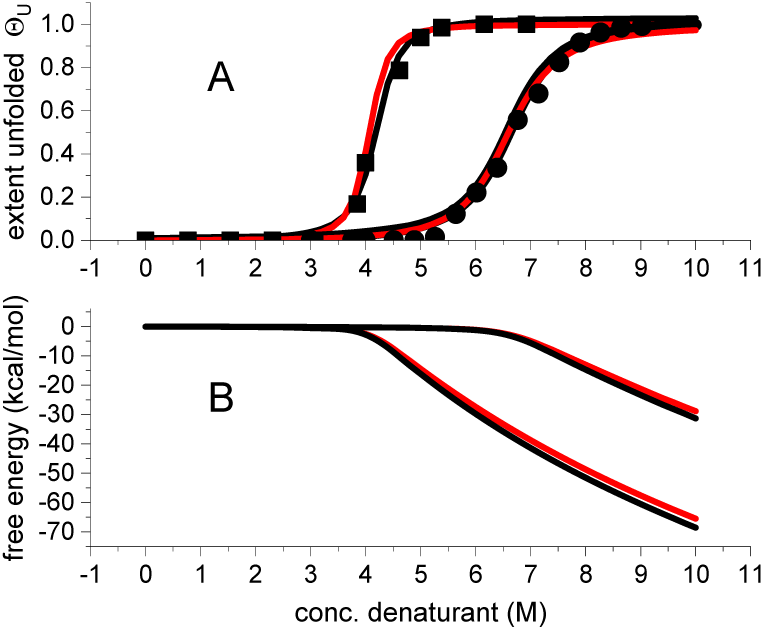
Chemical unfolding of lysozyme with guanidineHCL (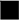) and urea (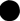) at 25 °C and pH 7. Data taken from reference.^16^ The midpoint concentrations c_0_ = 4.1 M for guanidine HCl cooperativity parameter is σ = 2×10^-3^ for both denaturants. The figure also compares the exact and approximate solutions for binding isotherm and free energy. (A) Extent of unfolding. Red line: matrix solution eq. 9. Black line: simplified isotherm eq. 17. (B) Free energy of unfolding. Red line: eq. 10. Black line: eq.18.

The urea-induced transition (unfolding concentration range Δc = 4.8 M) is broader than that of guanidineHCl (Δc = 3 M). However, in spite of these large differences, the corresponding diffusive or osmotic free energies ΔG_diff_ (eq. 20) constituted by these gradients are identical with ΔG_diff_ = −0.431 kcal/mol. Likewise, the free energy changes ΔF ≍ −24 kcal/mol and the cooperativity parameter σ = 2×10^-3^ are also identical for the two denaturants. The only significant thermodynamic difference is that the urea binding constant K_D_ = 0.15 M^-1^ is distinctly smaller than the guanidineHCl binding constant K_D_ = 0.245 M^-1^. The broadening of the transition region in urea solution is thus caused by the low urea binding constant and not by a change in the protein cooperativity.

Figure 3 compares the exact and the approximate solutions for the extent of unfolding and (eqs. 9 & 10 versus eqs. 17 & 18). The approximate solutions show an almost complete overlap with the exact solutions.

### Temperature-dependence of chemical unfolding of lysozyme

Lysozyme unfolding depends on pH and temperature. A decrease in pH or an increase in temperature shift the unfolding transition towards lower c_0_ values. This is illustrated in figure 4 for chemical denaturation with guanidineHCl at three different temperatures.

**Figure 4.**
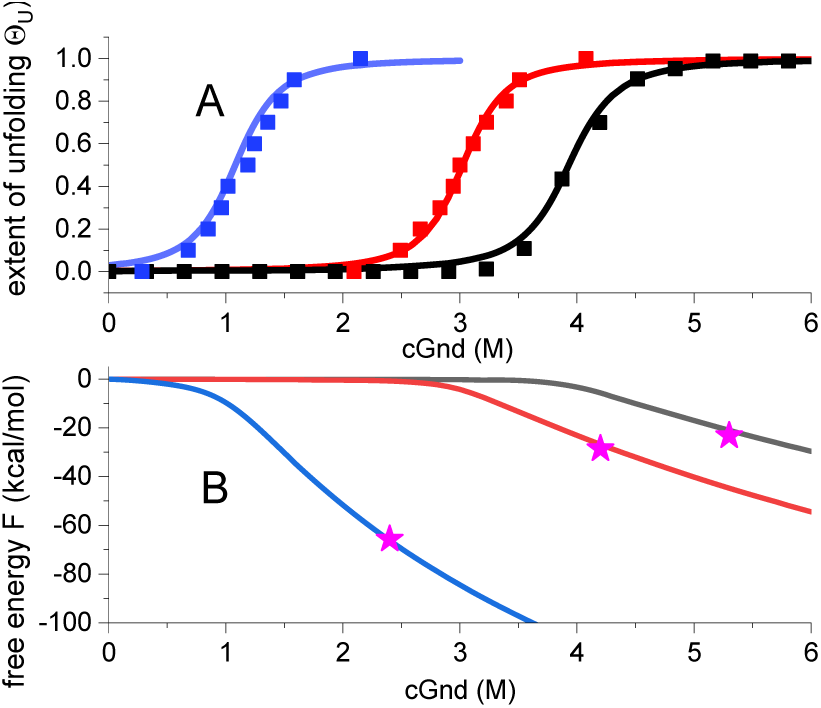
Temperature dependence of lysozyme unfolding in guanidineHCl solution. Data points taken from figure 2 of reference^17^. Solid lines are the simulations with the multistate cooperative model. Blue: 60 °C, K_D_ = 0.9 M^-1^, σ = 3×10^-2^. Red: 40 °C, K_D_ = 0.34 M^-1^, σ =2.5×10^-3^. Black: 10 °C, K_D_ = 0.26 M^-1^, σ = 1×10^-3^. (A) Extent of unfolding Θ_U_(c_D_). Simulations with eq. 9. (B) Free energy F(c_D_), calculated with eq. 10. (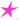 Free energies at 95% unfolding, calculated with eq. 19.

Figure 4A displays unfolding isotherms obtained with UV- and CD-spectroscopy.^17^ The solid lines are the simulations with the multistate cooperative model. The binding constants K_D_ are exactly the reciprocal of the midpoint concentrations c_0._ The binding constants increase with temperature and the transition regions broaden. The cooperative interactions are reduced, leading to a larger cooperativity parameter σ. A temperature increase from 10 °C to 60 °C increases the cooperativity parameter by a factor of 10.

Figure 4B displays the predicted concentration dependence of the free energy. The free energy is zero for the native protein and decreases rapidly beyond the midpoint of unfolding c_0_. The solid lines are calculated with the exact solution eq. 9. The three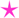-marked data points were calculated with eq. 19, corresponding to 95% unfolding. The comparison with the exact solution, demonstrate that the simple equation 19 is an excellent approximation for the free energy if c_D_ ≫c_0_.

We have analysed published denaturation experiments of lysozyme in guanidineHCl solutions at different temperatures and the results are summarized in table 1.

**Table 1.** Chemical unfolding of lysozyme with guanidineHCl at different temperatures

| Temp | C<sub>mid</sub> | K<sub>D</sub> | $\sigma$ | C<sub>ini</sub> | C<sub>end</sub> | $\Delta F$ | g<sub>den</sub> | $\Delta G_{H_2O}$ | m |
| --- | --- | --- | --- | --- | --- | --- | --- | --- | --- |
| °C | M |  |  | M | M | kcal/mol | kcal/mol | kcal/mol | kcal L/mol<sup>2</sup> |
| 10<sup>20</sup> | 3.8 | 0.263 | 1.50E-03 | 2.7 | 5.3 | -21.8 | -0.379 | 9.55 | 2.512 |
| 15<sup>20</sup> | 4.0 | 0.25 | 1.80E-03 | 2.7 | 5.6 | -22.7 | -0.417 | 8.598 | 2.150 |
| 20<sup>20</sup> | 3.9 | 0.257 | 2.00E-03 | 2.6 | 5.5 | -23.8 | -0.436 | 8.598 | 2.210 |
| 25<sup>20</sup> | 3.8 | 0.263 | 2.50E-03 | 2.4 | 5.5 | -26.1 | -0.491 | 9.554 | 2.513 |
| 35<sup>20</sup> | 3.3 | 0.303 | 5.00E-03 | 1.75 | 5.1 | -32.6 | -0.654 | 8.598 | 2.605 |
| 10<sup>17</sup> | 3.8 | 0.26 | 1.00E-03 | 2.9 | 5.25 | -20.1 | -0.334 | 9.55 | 2.483 |
| 40<sup>17</sup> | 2.9 | 0.34 | 2.50E-03 | 1.9 | 4.1 | -26.41 | -0.493 | 8.36 | 2.842 |
| 60<sup>17</sup> | 1.1 | 0.9 | 3.00E-02 | 0.25 | 2.4 | -65.5 | -1.5 | 4.299 | 3.869 |
| 25<sup>22</sup> | 3.9 | 0.256 | 1.00E-03 | 2.8 | 5.15 | -21.2 | -0.361 | 10.03 | 2.568 |
| 10<sup>21</sup> | 4.1 | 0.243 | 2.00E-03 | 2.7 | 5.8 | -22.55 | -0.43 | 8.837 | 2.147391 |
| 30<sup>15</sup> | 3.9 | 0.258 | 1.00E-03 | 2.9 | 5.3 | -21.7 | -0.363 | 8.837 | 2.236 |
| 45<sup>15</sup> | 3.2 | 0.31 | 1.00E-02 | 1.3 | 5.3 | -39.3 | -0.888 | 4.777 | 2.236 |

Polypeptides and proteins of different size and structure were also analyzed. The fits of the experimental data with the multistate cooperative model were all of comparable quality as shown for lysozyme in figures 2-4. The data are summarised in table 2.

**Table 2.** Chemical unfolding of peptides and proteins of different size and structure in guanidine Cl and urea

| Protein, Peptide | N<sub>aa</sub> | pH | Denaturant | K<sub>D</sub> | c<sub>mid</sub> | $\sigma$ | c<sub>ini</sub> | c<sub>end</sub> | $\Delta c$ | g<sub>den</sub> | $\Delta F$ | Lit. | Figure |
| --- | --- | --- | --- | --- | --- | --- | --- | --- | --- | --- | --- | --- | --- |
|  |  |  |  |  |  |  | M | M | M | kcal/mol | kcal/mol |  |  |
| BBL | 41 | 7 | GndHCl | 0.33 | 3.03 | 3.00E-02 | 0.2 | 7.5 | 7.3 | -2.14 | -20.9 | <sup>31</sup> | fig 2b |
| ubiquitin WT | 76 |  | GndHCl | 0.34 | 2.94 | 7.00E-03 | 1.4 | 5 | 3.6 | -0.753 | -22.1 | <sup>32</sup> | fig 1 |
| ubiquitin L67S | 76 |  | GndHCl | 0.72 | 1.39 | 1.30E-02 | 0.5 | 2.7 | 2.2 | -0.998 | -28.4 | <sup>32</sup> | fig 1 |
| ubiquitin | 76 | 2 | GnddHCl | 0.395 | 2.53 | 8.00E-03 | 1.2 | 4.5 | 3.3 | -0.782 | -24.1 | <sup>33</sup> | fig 4 |
| ubiquitin | 76 | 5.5 | GndHCl | 0.295 | 3.39 | 1.20E-02 | 1.3 | 6.3 | 5 | -0.934 | -26.4 | <sup>33</sup> |  |
| Apo lipoprotein A1 | 120 | 8 | GndHCl | 1 | 1.00 | 2.00E-02 | 0.3 | 2 | 1.7 | -1.12 | -49.7 | <sup>34</sup> | fig 1 |
| Apo lipoprotein A1 | 122 | 7.4 | GndHCl | 0.952 | 1.05 | 5.00E-03 | 0.6 | 1.7 | 1.1 | -0.61 | -32.8 | <sup>35</sup> | fig.7 |
| Apo lipoprotein A1 | 122 | 7.4 | GndHCl | 0.95 | 1.05 | 3.00E-02 | 0.3 | 2.3 | 2 | -1.21 | -55.9 | <sup>36</sup> |  |
| lysozyme | 129 | 7 | GndHCl | 0.245 | 4.08 | 2.00E-03 | 2.8 | 5.8 | 3 | -0.431 | -24.6 | <sup>16</sup> | fig 1 |
| lysozyme | 129 | 2 | GndHCl | 0.42 | 2.38 | 2.00E-02 | 0.8 | 4.6 | 3.8 | -1.03 | -49.6 | <sup>37</sup> | fig 2A |
| lysozyme | 129 | 2.5 | GndHCl | 0.48 | 2.08 | 5.00E-03 | 1.1 | 3.2 | 2.1 | -0.632 | -31.1 | <sup>37</sup> | fig 1 |
| PMS-Ct | 145 | 4 | GndHCl | 0.52 | 1.92 | 1.00E-03 | 1.1 | 2.5 | 1.4 | -0.486 | -22.7 | <sup>38</sup> | fig 4 |
| PMS-Ct | 241 | 4 | GndHCl | 0.513 | 1.95 | 4.00E-03 | 1.1 | 2.7 | 1.6 | -0.531 | -45.3 | <sup>38</sup> | fig 4 |
| **average** |  |  |  | **0.551** | **2.22** | **0.012** |  |  | **2.931** | **-0.90** |  |  |  |
| **STDV** |  |  |  | **0.256** | **0.93** | **0.0096** |  |  | **1.659** | **0.43** |  |  |  |
| Ac-tyr-(ala-glu-ala-ala-lys-ala)<sub>5</sub>-phe-NH<sub>2</sub> | 32 | 7 | urea | 0.25 | 4.00 | 8.00E-02 |  |  |  |  |  | <sup>6</sup> | fig 1 |
| Ac-tyr-(ala-glu-ala-ala-lys-ala)<sub>8</sub>-phe-NH<sub>2</sub> | 50 | 7.0 | urea | 0.2 | 5.00 | 1.00E-02 | 2.2 | 9.8 | 7.6 | -0.884 | -18.1 | <sup>39</sup> | fig. 1<br>273 K |
| HPr protein | 85 | 7 | urea | 0.215 | 4.65E+00 | 8.00E-03 | 2 | 8 | 6 | -0.793 | -26.2 | <sup>40</sup> | fig 1b<br>288 K |
| cytochrome c | 106 | 5 | urea | 0.2 | 5.00E+00 | 4.00E-03 | 2.9 | 7.7 | 4.8 | -0.578 | -25.1 | <sup>3</sup> |  |
| cytochrome c | 106 | 7 | urea | 0.155 | 6.46E+00 | 4.00E-03 | 3.6 | 9.5 | 5.9 | -0.574 | -22.5 | <sup>3</sup> | fig 5 |
| P22 1-domain | 123 | 6 | urea | 0.3 | 3.33 | 9.00E-03 | 1.5 | 5.6 | 4.1 | -0.753 | -35.18 | <sup>41</sup> | fig 2 |
| lysozyme | 129 | 2 | urea | 0.303 | 3.30 | 2.00E-02 | 2.2 | 6.4 | 4.2 | -0.632 | -47.9 | <sup>37</sup> | fig 2B |
| lysozyme | 129 | 5.5 | urea | 0.15 | 6.67 | 2.00E-03 | 4.5 | 9.3 | 4.8 | -0.43 | -23.3 | <sup>16</sup> | fig 1 |
| anti-EF-R mab | 1600 | 7 | urea | 0.18 | 5.56 | 2.00E-02 | 2 | 8.3 | 6.3 | -0.85 | -407 | <sup>3</sup> | fig 2 |
| **average** |  |  |  | **0.22** | **4.80** | **1.74E-02** |  |  | **5.46** | **-0.69** |  |  |  |
| **STDV** |  |  |  | **0.05** | **1.15** | **0.0229** |  |  | **1.12** | **0.15** |  |  |  |
| lysozyme | 129 |  | SDS | 230.00 | 0.00 | 7.00E-02 | 0.001 | 0.012 | 0.011 | -1.47 | -77.7 | <sup>18</sup> | Tab. 1<br>298 K |

### Chemical unfolding with sodium dodecyl sulfate (SDS)

Sodium dodecyl sulfate (SDS) is a much stronger denaturant then guanidine HCl or urea as shown in figure 5 (data taken from reference^18^). <u>U</u>nfolding was measured with calorimetry. At 25 °C the midpoint concentration of unfolding is only 4.35 mM, resulting in a large binding constant of K_D_ = 230 M^-1^. The cooperativity of SDS-induced unfolding is relatively low with σ = 7×10^-2^.

**Figure 5.**
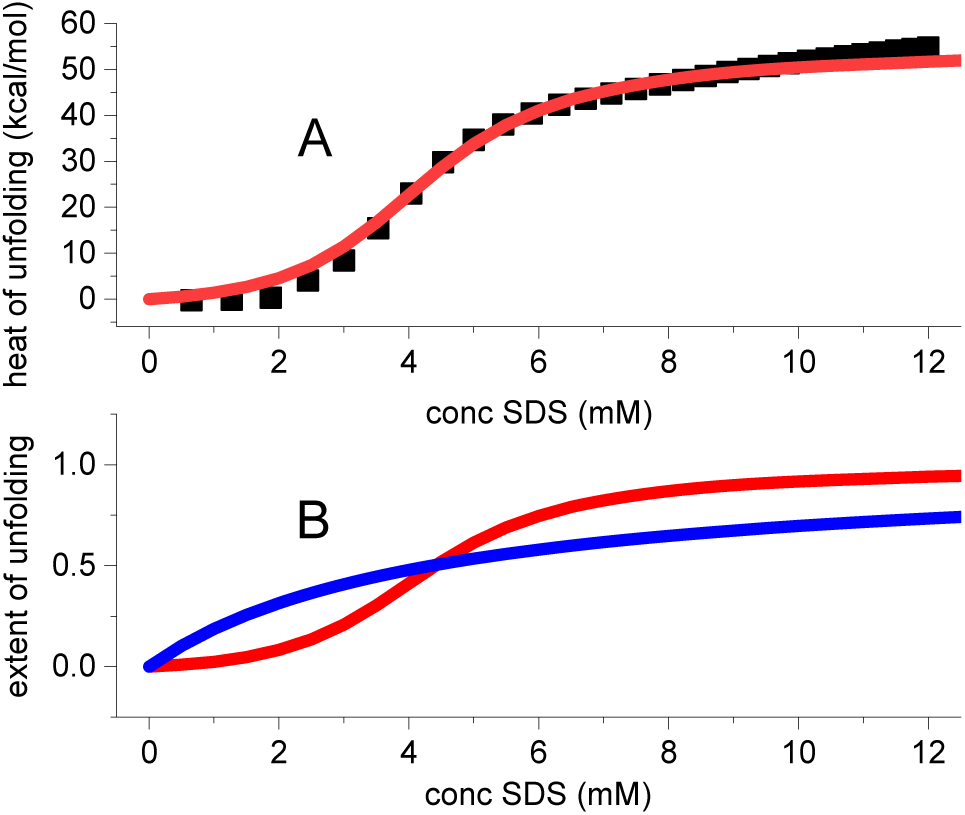
Calorimetric titration of 68 μM lysozyme with sodium dodecyl sulfate (SDS) solution at 25 °C. (A) Experimental data taken from reference,^18^ table 1. The published data were normalized to 1 mole lysozyme. Each data point is a separate measurement. Red line: multistate cooperative model. K_D_ = 230 M^-1^, σ = 7x 10^-2^, enthalpy of unfolding ΔH_0_ = 55 kcal/mol. (B) unfolding isotherm. Red line: multistate cooperative model, calculated with the same parameters as listed in panel A. Blue line: K_D_ as in panel A, but σ = 1 (non-cooperative Langmuir isotherm).

The enthalpy of unfolding, ΔH_0_, is the most important parameter in thermal unfolding studies. Spectroscopic measurements of chemical unfolding isotherms cannot provide this thermodynamic property. It is obtained, however, by a direct calorimetric measurement as shown in figure 5. Lysozyme is titrated with low concentrations of sodium dodecyl sulfate (SDS) and the heat of reaction is measured in a calorimeter. Each data point in figure 5 corresponds to an independent measurement.^18^ The total unfolding enthalpy is ΔH_n_ = 55 kcal/mol. This is in agreement with the result of an early isothermal enthalpimetric titration of lysozyme with guanidineHCl at pH 2.5 and 40 °C yielding 56 kcal/mol.^19^

Altogether 22 chemical unfolding isotherms of polypeptides and proteins of different size and structure were analysed. The fits of the experimental data with the multistate cooperative model were all excellent. The results are summarized in table 2.

## Discussion

### Characteristics of the multistate cooperative model

The multistate cooperative model is based on a statistical-mechanical partition function that contains two molecular parameters, the binding constant K_D_ = 1/c_0_ and the protein cooperativity σ. The cooperativity parameter determines the steepness of the unfolding transition (figure 1). The model takes into account the number ν of amino acid residues participating in the unfolding reaction and makes the following predictions. (i) The free energy of the n → u transition Δg_nu_ is identical to the free energy provided by the concentration gradient ΔG_diff_ of complete unfolding (eqs. 20 & 21). (ii) The free energy of the native protein is the reference state with zero free energy. This is the minimum free energy of the stable protein. Unfolding requires energy, which is stored in the unfolded protein and can be delivered if the unfolded protein returns to its ground state. (iii) A simple approximation can be given (eqs. 17 - 19) which fits the experimental data extremely well and is as easy to apply as the LEM.

### Temperature dependence of unfolding

The temperature dependence of the binding constant K_D_ and the cooperativity parameter σ together with that of the free energy is displayed in figure 6 for lysozyme (cf. Table 1). The simulation of the unfolding isotherms yields excellent fits in each case. Nevertheless, the scatter of the data in figure 6 is considerable as the isotherms are obtained by different authors under different experimental conditions. Two temperature regions can be discerned. Between 10 °C and 35 °C the effect of temperature is small, above 35 °C all properties change rapidly.

**Figure 6.**
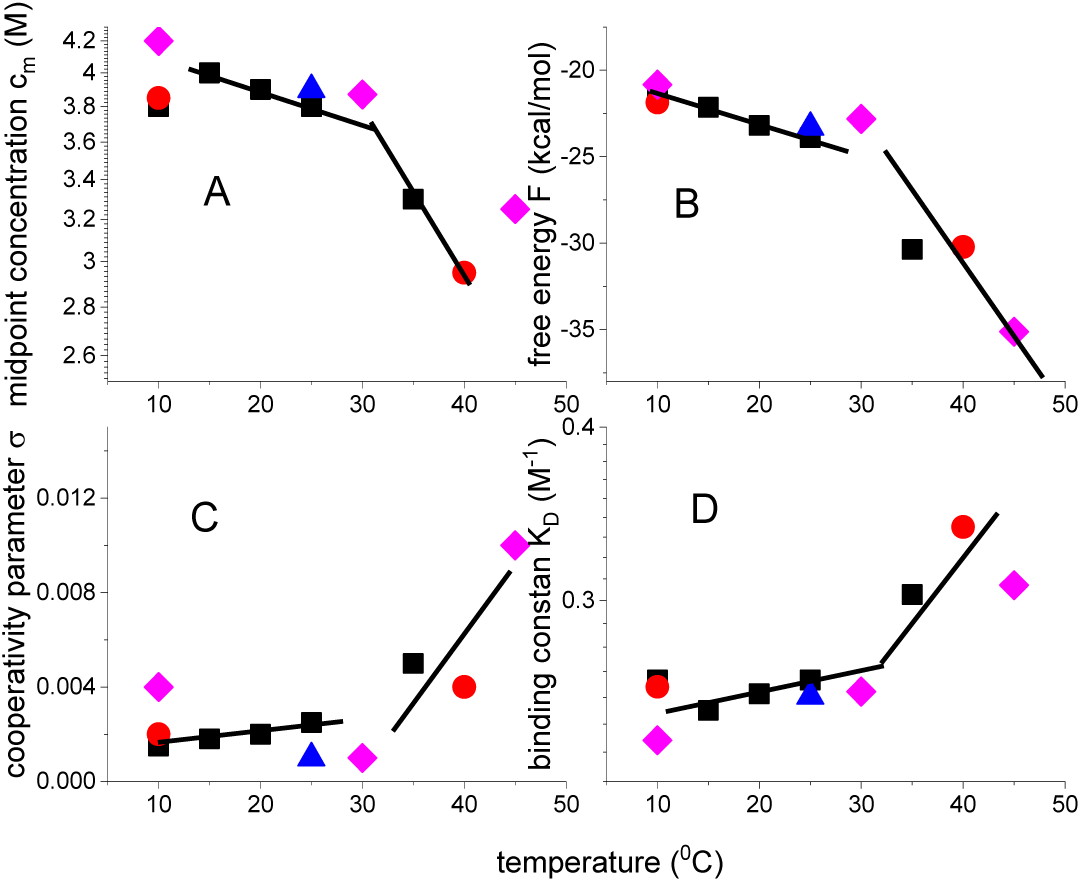
Lysozyme unfolding experiments in guanidineHCl at different temperatures and by different authors. (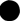)^20^, (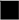)^17^, (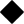)^15, 21^, (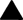)^22^. The solid lines are introduced to guide the eye.

The midpoint concentration and the free energy decrease with temperature, the cooperativity parameter and the binding constant increase. At higher temperature the protein cooperativity decreases. The cooperativity parameter of chemical unfolding is two orders of magnitude larger than that of terminal unfolding.^11, 23–25^ Chemical unfolding is less cooperative than thermal unfolding.

A detailed analysis of the temperature dependence of the free energy is given in figure 7. ΔH = ΔF + TΔS . Figure 7A displays data obtained from a consistent set of experiments for the temperature range 10 °C to 25 °C (figure 1 in reference^20^).

**Figure 7.**
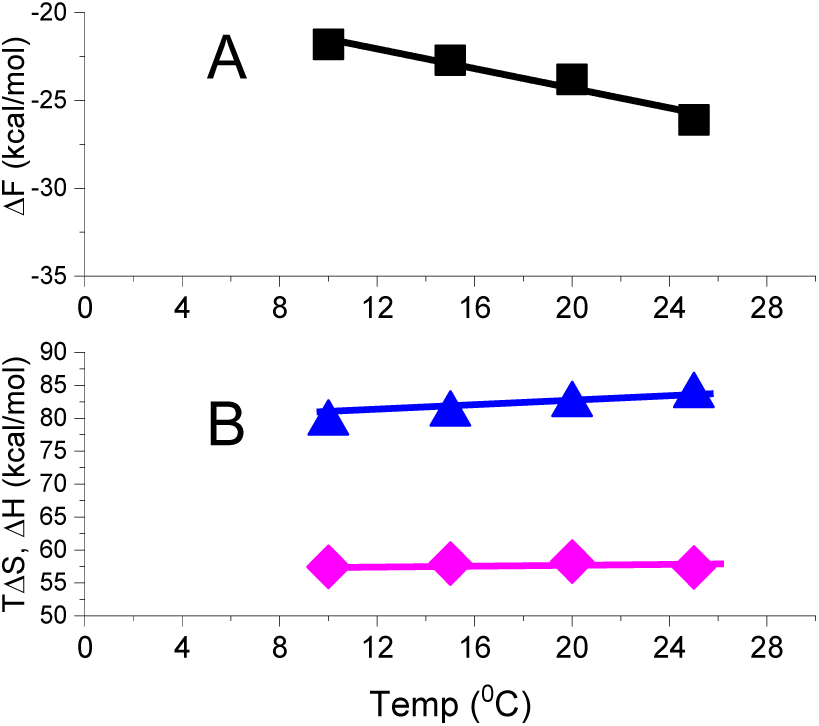
Chemical unfolding of lysozyme in guanidineHCl solution at different temperatures. (A) Free energy ΔF(T) calculated with the multistate cooperative model. Evaluation of unfolding isotherms of reference.^20^ Straight line: linear regression analysis with slope -ΔS = −0.28 ± 0.046 kcal/molK (R^2^=0.947). (B) (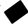) Enthalpy ΔH, (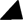) Entropy TΔS.

From the negative slope of the straight line in figure 7A the entropy 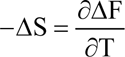 calculated as ΔS = 0.280±0.047 kcal/molK. Figure 7B then displays the entropy term TΔS and the predicted enthalpy ΔH = ΔF + TΔS . The average enthalpy is ΔH_0_ = 58 ± 1 kcal/mol. An early isothermal titration study of lysozyme with guanidineHCl, at pH 2.5 and 40 °C yielded ΔH = 56 kcal/mol.^26^. The present result is also in agreement with the SDS isothermal calorimetry shown in figure 5, resulting in an enthalpy of 55 kcal/mol at 25 °C

The general understanding in the protein folding field has been that proteins fold into their native conformations driven by a decrease in free energy (negative Δ*G*).^7^ The native protein is, thermodynamically, the most stable conformation. In the so-called funnel hypothesis, the native protein sits at the bottom of a rough-edged funnel representing the minimum free energy. The multistate cooperative model is fully consistent with this hypothesis. The native protein is the reference state with zero free energy and unfolding requires energy. In returning reversibly to the native state, the unfolded protein loses its free energy.

### Guanidine HCl and urea binding constants

Different molecular mechanism of denaturation have been proposed.^5, 27^ An indirect mechanism postulates changes in the water structure and of the hydrophobic effect. The alternative view is the direct interaction of the chemical denaturant with the protein. Strong support for the latter mechanism comes from isothermal titration calorimetry (ITC) of guanidineHCl and urea with various proteins.^28^ X-ray studies also demonstrate a close interaction between guanidineHCl and the protein backbone of lysozyme.^29^

The guanidineHCl binding constants deduced from the lysozyme isotherms are 0.2 M^-1^ ≤ K_D_ ≤ 0.9 M^-1^ in the temperature range 10 – 60 °C (table 1). Urea binds with a lower affinity of K_D_ = 0.15 M^-1^ (at 25 °C). Isothermal titration calorimetry was used to measure guanidineHCl and urea binding to different proteins.^28^ The data were analyzed by assuming a set of independent non-cooperative binding sites.^28^ The binding constants deduced for lysozyme were K_D_ ∼0.4 – 0.8 M^-1^ for guanidine HCl and 0.06 M^-1^for urea. This is in broad agreement with the results of the multistate cooperative model.

The binding constants of guanidineHCl and urea are small and the binding of a single guanidineHCl molecule is not sufficient to induce the n → u transition of an amino acid residue. Only the cooperative binding of many denaturants induces protein unfolding.

The adsorption isotherm (equation 15) is applicable also to denaturation with organic solvents. Lysozyme denaturation isotherms have been reported for ethanol and DMSO.^30^. The midpoint concentrations and corresponding binding constants are 5 M and 0.25 M ^-1^for ethanol and 7 M and 0.17 M^-1^ for DMSO. In contrast, anionic sodium dodecyl sulfate (SDS) has a high affinity for overall cationic lysozyme^18^. Based on figure 1 of reference^18^ the midpoint concentration is ∼4.3 mM and the binding constant K_D_ ∼230 M^-1^.

### Free energy of the N ⇆ U equilibrium

The multistate cooperative model provides a simple adsorption isotherm to analyse chemical unfolding (equation 15). The only unknown parameter is the cooperativity σ. The free energy follows from the partition function (equations 10 or 18). The free energy of the native protein is zero and the concentration profile is shown in figure 1. Addition of denaturant initially leads to only a small negative free energy. The free energy becomes distinctly more negative at concentrations near and above the midpoint concentration c_0_. For large denaturant concentrations c_D_ ≫ c_0_ . a simple approximation of ΔF(c_D_) ≍ −RT_0_ N ln(K_D_c_D_) (equation 19) is valid. Numerical values of the free energy change ΔF = ΔF(c_en d_) − ΔF(c_ini_), calculated with equation 10, are listed in table 1. The extent of unfolding in these calculations is 0.01 ≤ Θ_U_ ≤ 0.95 . It should also be noted that the shape of the free energy in chemical unfolding (figure 1) is identical to the free energy temperature profile in temperature-induced unfolding.^11^ However, the free energy change ΔF in chemical unfolding is about three fold higher more negative than the free energy of thermal unfolding.^11^ This must be traced back to the binding of the denaturants.

### Cooperativity in chemical and thermal unfolding

Chemical σ parameters are plotted as a function of temperature in figure 6. Cooperativity is σ ≃ 3 x 10^-3^ for temperature below 30 °C and strongly increases at higher temperature. The increase in temperature facilitates chemical unfolding by decreasing the cooperativity of the protein. The free energy to start a new folded sequence within an unfolded region is given by ΔF_σ_ = −RT ln σ.

## Conclusions

It is obvious that the Linear Extrapolation Method is a poor model to describe chemical unfolding. The two fit parameters have no well-defined thermodynamic or molecular basis. It is therefore attractive to modify an existing multistate cooperative model for the thermal protein unfolding to address chemical unfolding. The new two-parameter model, includes the binding constant of the denaturant and the cooperativity of protein unfolding. It is simple and easy to apply as the binding constant is the reciprocal of the midpoint concentration of unfolding. Simple equations can be given for the binding isotherm and the free energy. The free energy profile of chemical unfolding parallels the experimental results for thermal unfolding. The free energy of the native protein is zero and becomes distinctly negative upon unfolding, only after reaching the midpoint concentration. The new model is satisfying as it provides an excellent description of chemical denaturation isotherms and for the first time allows comparing cooperative, chemical and thermal unfolding.

